# Dye-based Fluorescent Organic Nanoparticles, New Promising Tools for Optogenetics

**DOI:** 10.1101/2024.06.10.598285

**Authors:** Jeremy Lesas, Thomas Bienvenu, Eleonore Kurek, Jean-Baptiste Verlhac, Delphine Girard, Frédéric Lanore, Mireille Blanchard-Desce, Cyril Herry, Jonathan Daniel, Cyril Dejean

## Abstract

Dye-based fluorescent organic nanoparticles are a specific class of nanoparticles obtained by nanoprecipitation in water of pure dyes only. While the photophysical and colloidal properties of the nanoparticles strongly depend on the nature of the aggregated dyes, their excellent brightness in the visible and in the near infrared make these nanoparticles a unique and versatile platform for in vivo application. This article examines the promising utilization of these nanoparticles for in vivo optogenetics applications. Their photophysical properties as well as their biocompatibility and their capacity to activate Chrimson opsin in vivo through fluorescence reabsorption process are demonstrated. Additionally, an illustrative example of employing these nanoparticles in fear reduction in mice through close-loop stimulation is presented. Through an optogenetic methodology, the nanoparticles demonstrate an ability to selectively manipulate neurons implicated in the fear response and diminish the latter. Dye-based fluorescent organic nanoparticles represent a promising and innovative strategy for optogenetic applications, holding substantial potential in the domain of translational neuroscience. This work paves the way for novel therapeutic modalities for neurological and neuropsychiatric disorders.

## 1 Introduction

Chronic electrical stimulation of brain areas is used in the treatment of many severe neuropsychiatric disorders[1, 2, 3, 4]. Although promised to become an alternative treatment of choice in several neuropsychiatric disorders, as it is already for Parkinson’s disease, portable brain modulation devices remain of limited use, mainly because of their inconsistent efficacy. It is likely that inconsistent outcomes result from their inability to precisely manipulate defined functional neuronal networks. With much room for improvement, new exploratory brain stimulation trials could be enabled by an increase in spatial and temporal specificity towards functional efficiency. Today, optogenetics is one of the greatest therapeutic hopes for brain stimulation because it allows for the focal manipulation of targeted neural circuits with great temporal precision. Its principle is based on the use of a light source to selectively modulate the activity of neurons transduced to express photosensitive ion channels (opsins). The spatio-temporal specificity of optogenetics made the technique a breakthrough in the possibility to target specific neuronal mechanisms, limiting side effects due to off-target stimulation. For instance, in preclinical animal models, recent work showed that targeted close-loop optogenetic manipulation of neuronal activity is a mean to reduce learnt fear expression in mice[5] or alleviate epileptic seizures[6]. Such close-loop optogenetic strategies could enhance efficiency in treating many psychiatric (major depression, anxiety disorders, obsessive-compulsive disorder) and neurological disorders such as epilepsy or chronic pain,[7, 8] for which first line pharmaco-logical therapies can become uneffective or inadequate.

Current optogenetic techniques require the delivery of light to the target brain region or organ at wavelength in the visible spectrum. While opsins can be activated by visible light only, this range of wavelength is highly scattered and attenuated in brain, requiring the insertion of optical fibers directed at the target. There are two major issues inherent to that approach. First, such procedures come with various hazards for the subject, such as direct damage to the tissue, in situ deterioration of the fiber and foreign object immune response. Second, the stimulated area is only as large as the cone of light of the optic fiber, that has to remain of small diameter for aforementioned safety reasons. Hence, optogenetic cannot be applied to volumes as large as human potential cerebral targets. All in all current approaches hinders the transposition of optogenetics to clinics. To increase the efficiency and topographical reach of optogenetics an innovative methodological angle has been proposed in the addition of fluorescent nanoparticles used to circumvent light scattering in the tissue. Designed nanoparticles have been assembled with compounds able to absorb and emit photons at controlled wavelength and in a radial manner with further implemention for neuronal stimulation in rodents[9, 10]. While this approach is very effective, the inorganic compounds used so far have limited therapeutic potential as they present limitation in both colloidal stability and brightness and can make use of elements suspected to be non biocompatible following long-term decaying processes[11]. Hence while optogenetics-nanoparticles combinations bear hopes for precision therapeutics, innovative improvements are needed to reach clinical standards, namely i) efficacy, ii) stability and iii) biocompatibility.

In that perspective, fluorescent organic nanoparticles (FONs) for brain stimulation represent a desirable alternative to inorganic instances. These nanoparticles are only composed of molecules or macromolecules and can hold bright fluorescence properties in water[12] while maintaining colloidal stability. Among the broad family of FONs, the most representatives are i) nanoparticles made of polymers - e.g. doped with commercial fluorescent dyes[13] or synthesized using fluorescent co-block polymers[14]- and ii) AIE nanoparticles (Aggregation Induced Emission) - which consist in restoring the fluorescence properties of non fluorescent dyes upon aggregation (mostly exploiting the restriction of rotation of *π* conjugated moities through C-Si and C-N bonds, or E/Z isomerization)[15]-.

Among FONs, the class of dye-based Fluorescent Organic Nanoparticles (dFONs) offer singular optogenetic-prone properties [16, 17]. These noncrystalline dye-based nanoparticles are prepared by nanoprecipitation of pure hydrophobic polar and polarisable fluorescent dyes in aqueous solution. They are self-stabilized (i.e. do not require the use of surfactants nor solubilizing coating) and bear intense brightness properties competitive with the brightest inorganic nanoparticles (e.g. Quantum Dots)[18] or fluorescent commercial dyes[19]. Unlike AIE, the aggregation of fluorescent dyes into dFONs induces a loss in fluorescence efficiency (i.e. Aggregation Caused Quenching or ACQ)[17] which strongly depends on the dipole-dipole interactions between aggregated dyes[20]. This loss in fluorescence can be limited through dedicated molecular design like the use of propeller shape dyes,[21, 17] or the addition of bulky groups[17, 19]. Note that dFONs are prepared with a minimal amount of water-miscible volatile organic solvent (*<* 1% for dFONs whereas it can reach *>*10% for AIE),[15] limiting the potential toxicity of the aqueous suspension to the sole nanoparticles. The specificity of dFONs lies in the nature of their constitutive polar and polarizable dyes, which governs i) the photophysical properties of dFONs and ii) ensures their colloidal stability. The brightness of the dFONs is also highly dependent on the overall size of the nanoparticle (i.e. the number of nano-confined dyes). As such, increasing the volume of the nanoparticles will result in brighter dFONs without changing their absorption and emission spectra. In addition to be efficient bright nanoparticles [12], dFONs can be optimized to give access to excitation in optical windows of longer wavelength [17], which are the more suitable for travelling in brain tissue and therefore modulate larger volumes.

*The aim of this work is to combine the outlaid approaches and develop an expandable toolbox for dFONs use in biological applications, with ambitions for* l*ong-term transposition into clinical opportunities. After their assembly and characterization we successfully used dFONs in combination with optogenetics in mice to manipulate neuronal activity in vivo without notable insults to the tissue. Strong of this proof of concept, we then achieved the manipulation of a brain function, fear expression, through a temporally and spatially focused close-loop stimulation strategy. Our study demonstrate that the formulation of hydrophobic dyes is an effective strategy for in vivo application of photoconversion, in our case for optogenetic stimulation mediated by fluorescence reabsorption. Our study paves the way for a standard development of versatile dFONs toolboxes with various absorption-emission spectra, with hopes for clinical use*.

## 2 Results and Discussion

### 2.1 Design of dFONS

The strategy we developed to activate opsins relies on the direct re-absorption of the fluorescence of nearby dFONs. Based on our previous work, we selected **A** as a dye sub-unit for dFONs because its fluorescence spectrum have a favourable overlap with the absorption spectra of the Chrimson opsin [22]. Plus, we demonstrated that dFONs **A** can be excited both in the visible - by the classical excitation pathway- and in the NIR-I exploiting their high two-photon absorption properties 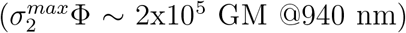 [23], giving access to deeper excitation in tissues. Moreover, we anticipated that this spectral compatibility allows for the excitation through space of the opsin by direct reabsorption of the fluorescence of dFONs **A**. Yet, despite brightness with orders of magnitudes higher than endogenous (e.g. flavine mononucleotide: 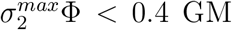 [24] or fluorescent proteins (e.g. tdTomato: ∈^*max*^Φ=3×10^5^m^−1^.cm^−1^ and 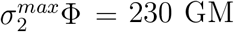 [24, 25], the efficiency of opsin activation by dFONs **A** could be affected by the extremely low concentration of dFONs (stock suspension ~ nM range). Then depending on the fate of the dFONs in vivo (i.e. diffusion, immunogenic response, degradation, etc…), their concentration may decrease dramatically, reducing the number of emitting dFONs, and consequently, the probability of exciting neurons expressing Chrimson. Moreover, the use of an excessively high concentration of dFONs (e.g. µm range) may raise concerns about colloidal stability of the dFONs as well as immune responses. Taking this into account, we set to optimize the brightness of dFONs rather than increase their concentration.

In order to maximize brightness, we designed bicomponent nanoparticles, where dFONs **A** are layered with another dye: such layering is named core-shell. These specific conditions, i.e dyes are polar and polarisable with similar chemical structure and good spectral overlap, deeply modify the photophysical properties of the core[26, 27]. The close proximity of core-layer and shell-layer dyes at the nanointerface of the core-shell structure promotes energy transfers between the two entities, leading to a vanishing fluorescence of the shell dyes accompanied with a pronounced enhancement in the fluorescence of the core dyes. Another characteristic of this phenomenon known as Nano-Interface Enhanced Emission (NIEE) manifests as subtle shifts in the fluorescence of core dyes towards shorter wavelengths[26]. In the case of dipolar triphenylamine-based push-pull dyes with fused-thiophene bridge, a 20 times enhancement of the fluorescence quantum yield has been reported[27]. To harness the NIEE, dye **D** emerges as a promising candidate for core-shell like dFONs in conjunction with **A** as i) both dyes have similar chemical structure, ii) the emission spectrum of fluorescent dye **D** overlaps well with the absorption of **A**, and iii) **D** maintains its fluorescence properties when aggregated (table 1), which is a key feature for efficient energy transfer. Moreover, the use of **D** to layer dFONs **A** is expected to prevent the potential degradation of **A** caused by water, particularly through retro-Knoevenagel reaction, thereby ensuring the preservation of core **A**’s fluorescence properties that is essential for long term use. The bicomponent dFONs **A**@**D** were prepared by a two-step procedure using the so-called nanoprecipitation method (Figure 1) of hydrophobic dyes **A** and **D** in pure water. First, dFONs **A** were prepared in water, followed by the gradual addition of **D** to encourage the seeding of **D** onto dFONs **A**, rather than the formation of standalone dFONs **D**. The resulting was a clear and coloured solution that both evidences the successful suspension of dyes in water and suggests the presence of distinct nanoparticles in water without evidence of large aggregates. Through their brightness and overall stability, these core-shell nanoparticles offer a strong compatibility potential for biological use. In comparison, other approaches such as expanding the amount of dye per nanoparticle or decorating the surface of dFONs with polymers [28] are problematic in biological media since they lead to larger nanoparticles, or pertain long-term stability of colloidal properties and brightness in biological serum. Stemming from **A**@**D** dFONS favorable biocompatibility potential, we set to further characterize them and implement their use in a biological system.

**Table 1:** Photophysical properties of the dyes subunits in water and the dFONs in water.

| Sample | $\emptyset^{(1)}$<br>nm | $\lambda_{abs}^{max}$<br>nm | $\epsilon^{max,2)}$<br>$M^{-1}.cm^{-1}$ | $\epsilon\Phi_f^{max,2)}$<br>nm | $\lambda_{em}^{max}$<br>$M^{-1}.cm^{-1}$ | $\Phi_f^{(3)}$ | $\lambda_{2Pabs}^{max}$<br>nm | $\sigma_2^{max,2)}$<br>GM | $\sigma_2\Phi_f^{max,2)}$<br>GM |
| --- | --- | --- | --- | --- | --- | --- | --- | --- | --- |
| **D** | — | 378 | $3.8 * 10^4$ | $3.8 * 10^3$ | 509 | 0.10 | 720 | 150 | 15 |
| **A** | — | 450 | $4.5 * 10^4$ | $0.6 * 10^3$ | 644 | 0.01 | 720<br>910 | 210<br>170 | 3<br>2 |
| **A@D** | — | 450 | $4.4 * 10^4$ | $1.4 * 10^3$ | 600 | 0.03 | 730<br>890 | 360<br>170 | 12<br>6 |
| dFONs **A** | 45 | 450 | $1.6 * 10^8$ | $2.2 * 10^6$ | 644 | 0.01 | 720<br>910 | $7.8 * 10^6$<br>$6.3 * 10^6$ | $1.1 * 10^5$<br>$0.9 * 10^5$ |
| dFONs **A@D** | 44 | 450 | $1.6 * 10^6$ | $5.3 * 10^6$ | 600 | 0.03 | 730<br>890 | $13.3 * 10^6$<br>$6.3 * 10^6$ | $4.4 * 10^5$<br>$2.0 * 10^5$ |

<sup>1)</sup> Dry diameter measured by transmission electronic microscopy.
<sup>2)</sup> For dFONs: calculated using size and density relative to water (d= 1.0, number of dyes per dFON= 37000).
<sup>3)</sup> Fluorescence quantum yield measured using fluorescein in 0.1 M NaOH<sub>aq.</sub> as standard ( $\Phi_f = 0.90$ ).

**Figure 1:**
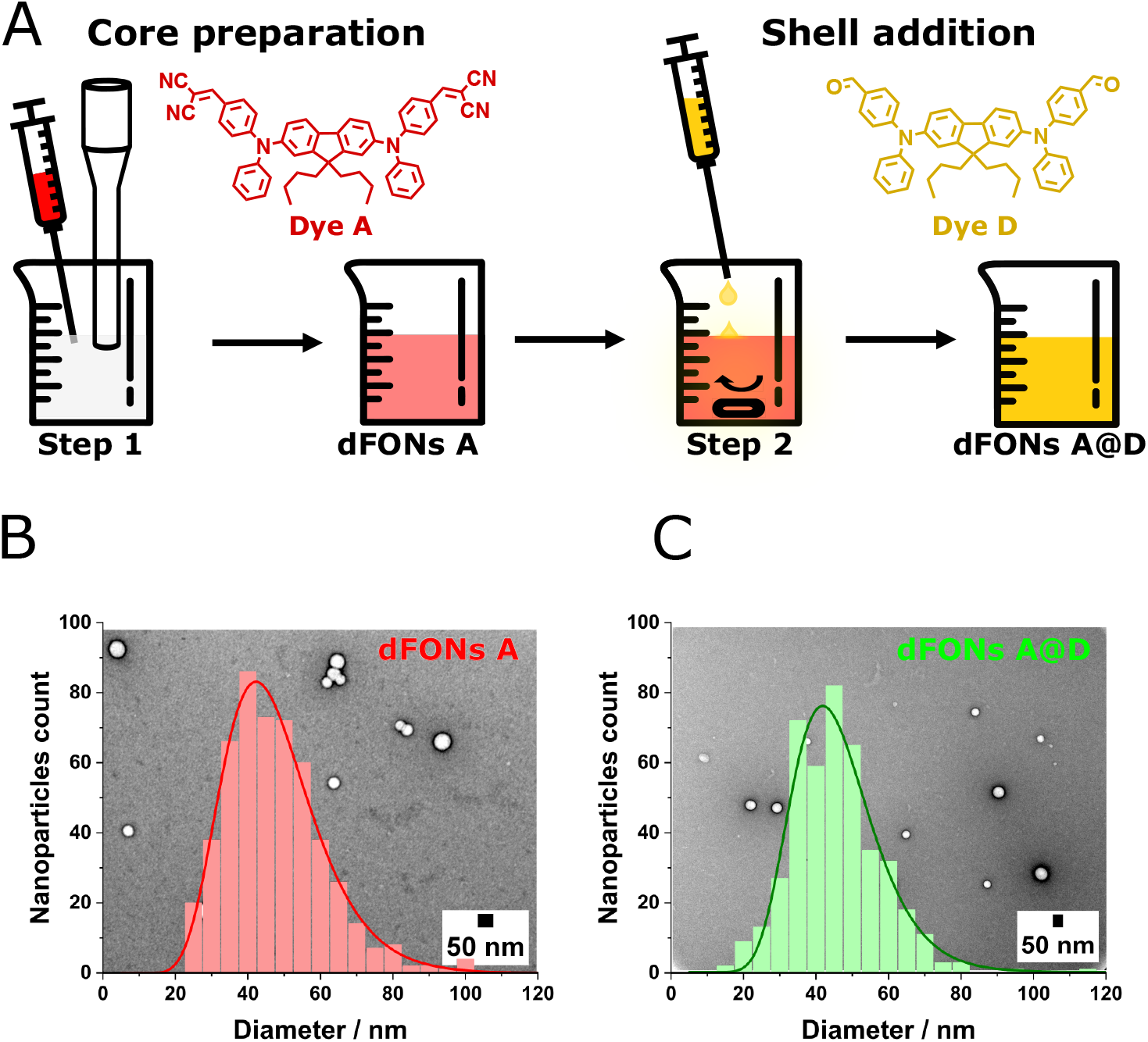
Preparation and size of dFONs. Preparation steps of **A@D** dFONs (A). Size distribution (with log normal fitting curve) and image of transmission electronic microscopy of dFONs **A** (B) and dFONs **A**@**D** (C).

### 2.2. Characterization of dFONs

#### 2.2.1 Size properties of dFONs

Electronic microscopy confirmed the presence of small nanoparticles in dFONs **A** and dFONs **A**@**D** samples, with mean dry diameters of 45 and 44 nm respectively (Figure 1). Interestingly, we found that the diameter of dFONs **A** was superior by 25% compared to what we reported earlier[23]. This difference can be explained by the use of a sonication probe (instead of a slow magnetic stirring) in the preparation of nanoparticles which led to a different size and possibly a change in molecular packing within the nanoparticles, as suggested by electronic molecular spectroscopy (see photophysical properties of dFONs in water section). TEM histograms revealed a single population for both dFONs with a broad dispersion in size (FWHM ~ 30 nm) typical of nanoparticles obtained by nanoprecipitation. Strikingly, the size distribution as well as the mean diameter of the bicomponent dFONs **A**@**D** are very close to the ones of pure dFONs **A** (Figure 1). Considering the expected quantity of dye to be layered onto the dFONs **A**, we anticipated, through calculation, an increase in size distribution of approximately 5-6 nm (assuming uniform packing density for both dyes). It is noteworthy that this disparity likely takes place during the layering process: as a solution of **D** in THF is added dropwise, there sould be competitions between i) the seeding of **D** - forming dFONs **D**-, ii) the layering of dFONs **A** - forming dFONs **A**@**D**- and iii) the dissolution of the surface of dFONs **A** - due to the presence of a gradient concentration of THF, depending on the mixing conditions and diffusion coefficients of water and THF[29]-. We hypothesize that the mild dissolution of the surface of the dFONs **A** (due to the good solubility of **A** in THF) is the key parameter that rules the size of the resulting dFONs **A**@**D**.

#### 2.2.2 Photophysical properties of dFONs in water

Absorption of aqueous dFONs **A**@**D** shows a large and intense band in the visible range (i.e. 400-550 nm) with a clear positive offset of the absorption band in the UV regarding the absorption spectrum of standalone dFONs **A** (Figure 2). Interestingly, we observed an overlay between the absorption spectrum of dFONs **A**@**D** and a “reference” absorption spectrum obtained by mixing the two dFONs **A** and **D** (i.e. a solution with independently prepared dFONs **A** and dFONs **D**, 1:1, v/v). This overlay confirms that the hyperchromic effect observed in the UV is mostly due to the absorption of **D**. This also demonstrates the presence of **D** (at the expected concentration) in the dFONs **A**@**D** dispersion (Figure 2).

**Figure 2:**
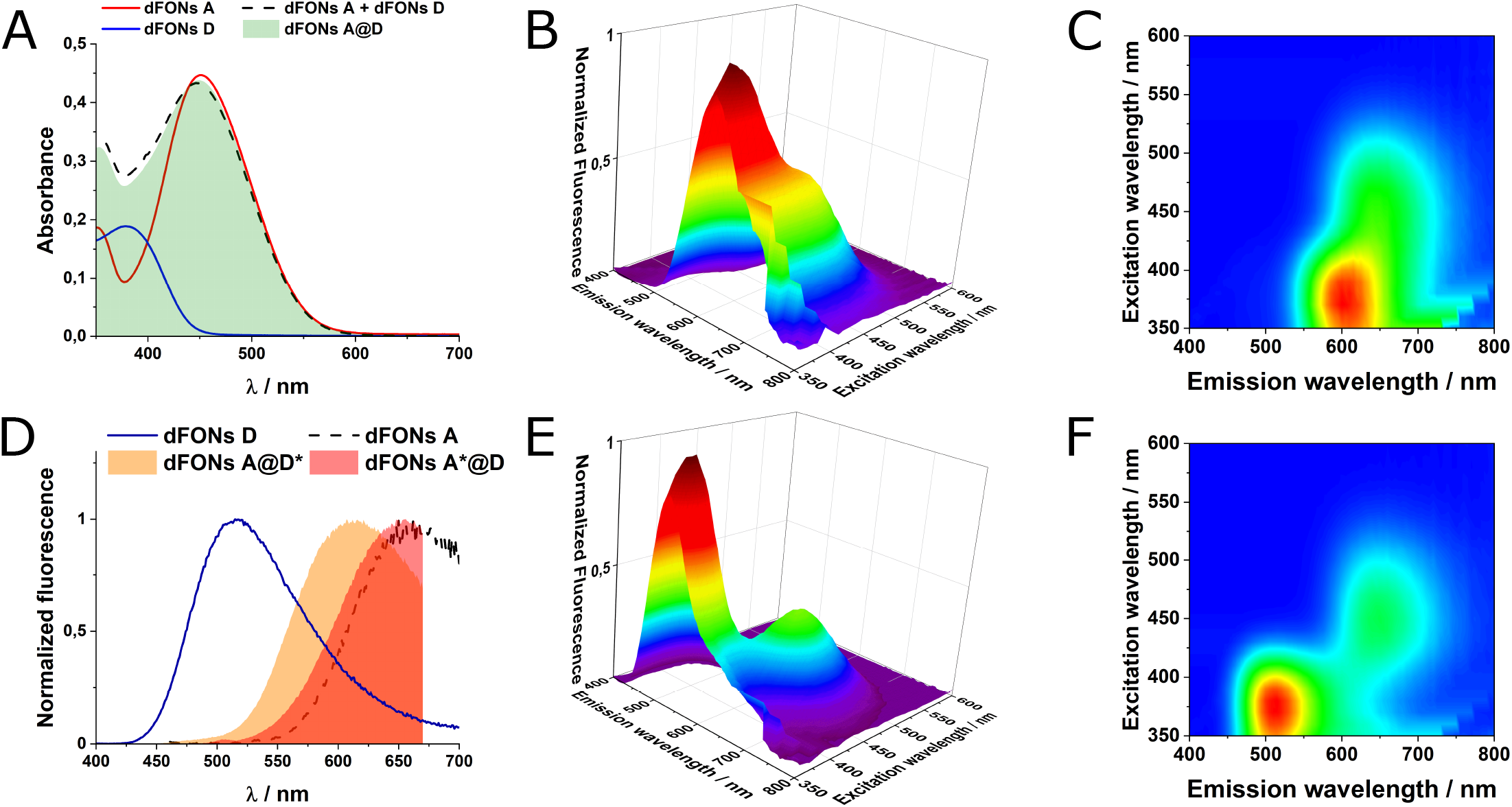
One-photon absorption properties of dFONs in water. Absorption (A) and normalized emission (D) spectra of dFONs. Native absorption of “dFONs **A** + dFONs **D**” has been corrected with dilution factor of 2. For the emission of dFONs **A**@**D**, fluorescence depends on the direct (**A**\*: *λ*^*ex*^= 450 nm) or indirect (**D**\*: *λ*^*ex*^= 380 nm) excitation. 3D excitation-emission measurements on dFONs **A**@**D** (B, C) and a solution of dFONs **A** with dFONs **D** in a ratio 1:1 (v/v, E, F). B and E represent the full 3D plot while C and F are the projection of the corresponding 3D plot on the fluorescence intensity axis.

Note that the presence of **D** does not prove that the dye has been precipitated over dFONs **A** to form bicomponent dFONs **A**@**D**. Yet, when exciting preferentially **D** (*λ*^*ex*^= 380 nm), the emission of dFONs **A**@**D** reveals the presence of a single red fluorescence band (*λ*^*em*^ = 500-700 nm) peaking at 600 nm (Figure 2) without any evidence of fluorescence from **D**. While the emission range is closer to that of dFONs **A**, the fluorescence process (i.e. Φ_*f*_) of the bicomponent dFONs is much more probable, with a three-fold increase (Table 1). We interpret such modifications in the de-excitation (vanishing of dFONs **D** emission) and emission properties (enhancement of dFONs **A** fluorescence) as the direct consequence of energy transfer from dyes **D** to **A** in dFONs **A**@**D** thanks to NIEE effect.

To validate this interpretation, we measured 3D-excitation/emission spectra of i) dFONs **A**@**D** and ii) a mixed solution containing both pure dFONs **A** and pure dFONs **D** (1:1, V/V). As evidenced on Figure 2, red fluorescence of dFONs **A**@**D** does not come from energy exchange or reabsorption process between dFONs **D** and dFONs **A**. Moreover, excitation and emission spectra show that there are indeed 2 emitting centers: the most intense fluorescence comes from the energy transfer from **D** to **A** (*λ*^*ex*^ = 380 nm, *λ*^*em*^= 600 nm) - when exciting **D**- and the second one (*λ*^*ex*^ = 450 nm, *λ*^*em*^= 644 nm) comes from direct excitation of **A**. We hypothesized that this “double” emission is a direct consequence of NIEE effect; as we previously observed a marked blue shift of emission for core-shell nanoparticles built from highly polarizable push-pull dyes[26]. This phenomenon was attributed to the confinement of the energy transfer and emission at the interfacial layer between core and shell dyes. On the other hand, direct excitation of dFONs **A** (*λ*^*ex*^ *>* 450 nm) led to the excitation of all dyes **A** (i.e. not only those at the nanointerface with **D**). This collective response is therefore closer to the one of pure dFONs **A** (table 1). These results give clues to decipher the real composition of dFONs **A**@**D** and confirm a layering of **A** by **D**, thus, confirming the core-shell architecture.

Taking advantage of the enhancement of the fluorescence quantum yield by NIEE, we performed two-photon absorption (2PA) measurements using the two-photon excited fluorescence technic on dFONs in water (Figure 3). 2PA revealed that dFONs **A**@**D** absorb on a broad spectral domain in the NIR (*<* 700 nm up to *>* 1.00 µm) with its maximum at 730 nm and two sub-bands at 800 nm and 900 nm. As shown on Figure 3, the shape of the 2PA of dFONs **A**@**D** is close to dFONs **A** (same maximum and sub-band position). Yet the main absorption band in the range 700-800 nm is somewhat distended compared to dFONs **A**. Indeed, while the two-photon cross section (*σ*_2_) from 800 nm to 1.0 µm are the same for both dFONs **A**@**D** and **A**, dFONs **A**@**D** is a much better two-photon absorber below 800 nm (~ two folds).

**Figure 3:**
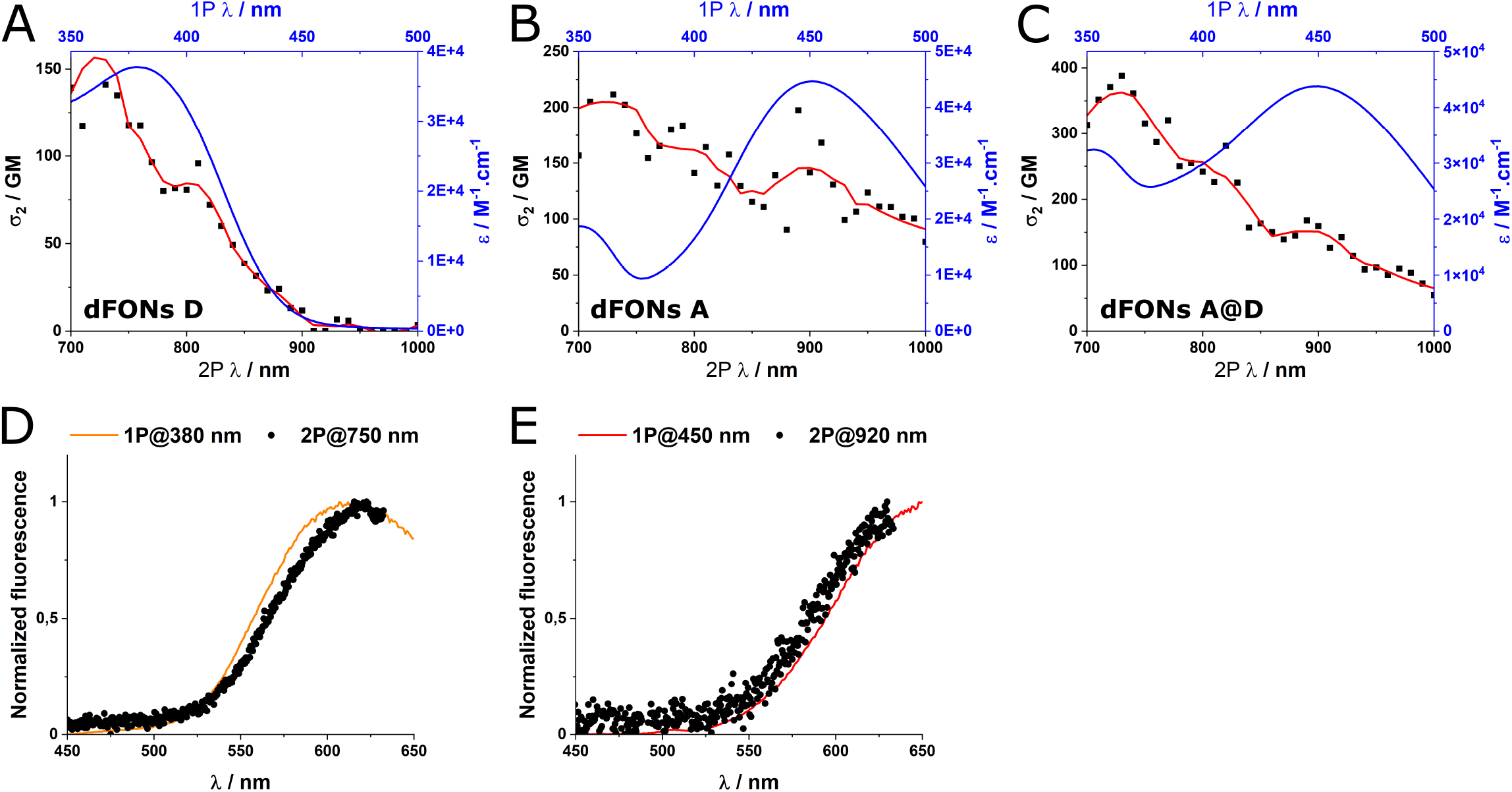
Two-photon properties of dFONs in water. Overlay of the two-photon and one-photon absorption spectra of dyes in dFONs **D** (A), **A** (B) and **A**@**D** (C) in water. Smooth functions are plotted as red lines to support reading of experimental data. Overlay of emission spectra of dFONs **A**@**D** measured by one- and two-photon excitation of **D** (D) or **A** (E). Note that full spectra are not available due to the presence of a shortpass filter to remove laser signal.

It is worth noting that the use of sonication for the preparation of dFONs **A** not only affects the size of the nanoparticles but also the dipole-dipole interactions. Indeed, while the absorption and emission maximums are slightly different (reported values, 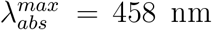, and 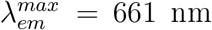)[23], the one and two-photon absorption are twice lower. This suggests that **A** adopts a configuration where the two dipoles are organized in an anti-parallel configuration, inducing an anti-cooperative effect, thus lowering the one and two-photon absorption. Remarkably, the two-photon absorption (2PA) spectrum of dFONs **D** exhibits a pronounced band below 800 nm (Figure 3), aligning well with the enhancement in the 2PA response observed with dFONs **A**@**D**. This alignment is consistent with the single-photon excitation spectrum presented in Figure 2. Further examination of the emission spectra, measured under two-photon excitation conditions (Figure 3D-E), revealed 2P bands where **D** strongly absorbs (e.g. *λ*^*ex*^= 750 nm) or only A absorbs (e.g. *λ*^*ex*^ *>* 920 nm). Intriguingly, the emission profiles perfectly coincide with the Nano-Interface Enhanced Emission (NIEE) at *λ*^*ex*^= 750 nm and the emission from pure **A** at *λ*^*ex*^= 920 nm.

Considering that the efficiency of energy transfer depends on the sixth power of the distance (Förster) or the probability of orbital overlap (Dexter) between the donor and acceptor (here, **D** and **A**), and taking into account the extreme dilution of dFONs (~ nM range) with the low probability of a two-photon process,the occurrence of emission resulting from energy transfer between two nanoparticles (**A** and **D**), obtained by two-photon excitation, is at least unlikely if not impossible. This demonstrates that dFONs **A**@**D** constitute core-shell structures with highly efficient energy transfer from dye **D** to **A** (full vanishing of **D** fluorescence), resulting in intensely bright red fluorescent organic nanoparticles excitable in the visible (*ϵ*^*max*^Φ = 5×10^6^ M^*−*1^.cm^*−*1^) and in the NIR-I 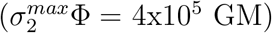. These characteristics position dFONs **A**@**D** nanoparticles as promising candidates for an efficient combination with optogenetics in vivo.

### 2.3 In vivo experiments

#### 2.3.1 Experimental strategy for in vivo demonstration of dFONs use

To assess the potency of dFONs for in vivo applications, we studied their efficacy in mediating optogenetic activation of neocortical neurons. The optogenetic strategy consisted in using a viral vector in order to infect neurons with engineered DNA coding for a light sensistive protein, an opsin. Opsin expressing neurons become sensitive to light, in a way that it is possible to manipulate their activity with specific wavelengths. Our preparation consisted in getting neocortical neurons to express both the Chrimson opsin which is sensitive to yellow-orange wavelengths and the tdTomato red fluorescent protein, serving as a visual reporter to confirm Chrimson expression (Figure 4). Chrimson opsin absorption band overlaps well with the emission spectra of dFONs **A@D**. In addition it has a low absorptivity in the UV (i.e. 365-405 nm range) allowing for the selective excitation of dFONs **A@D** (UV or NIR-I) and Chrimson. Hence, we expect neurons to be exctited by UV light only in the presence of dFONs and Chrimson. Since Chrimson spectrum is close to that of dFONs but also distant from their excitation maximum, the positive control condition will consist in delivering yellow light to directly activate the opsin, together with two negative control conditions consisting in the delivery of UV or yellow light i) without the dFONs and ii) without both the dFONs and the opsin.

**Figure 4:**
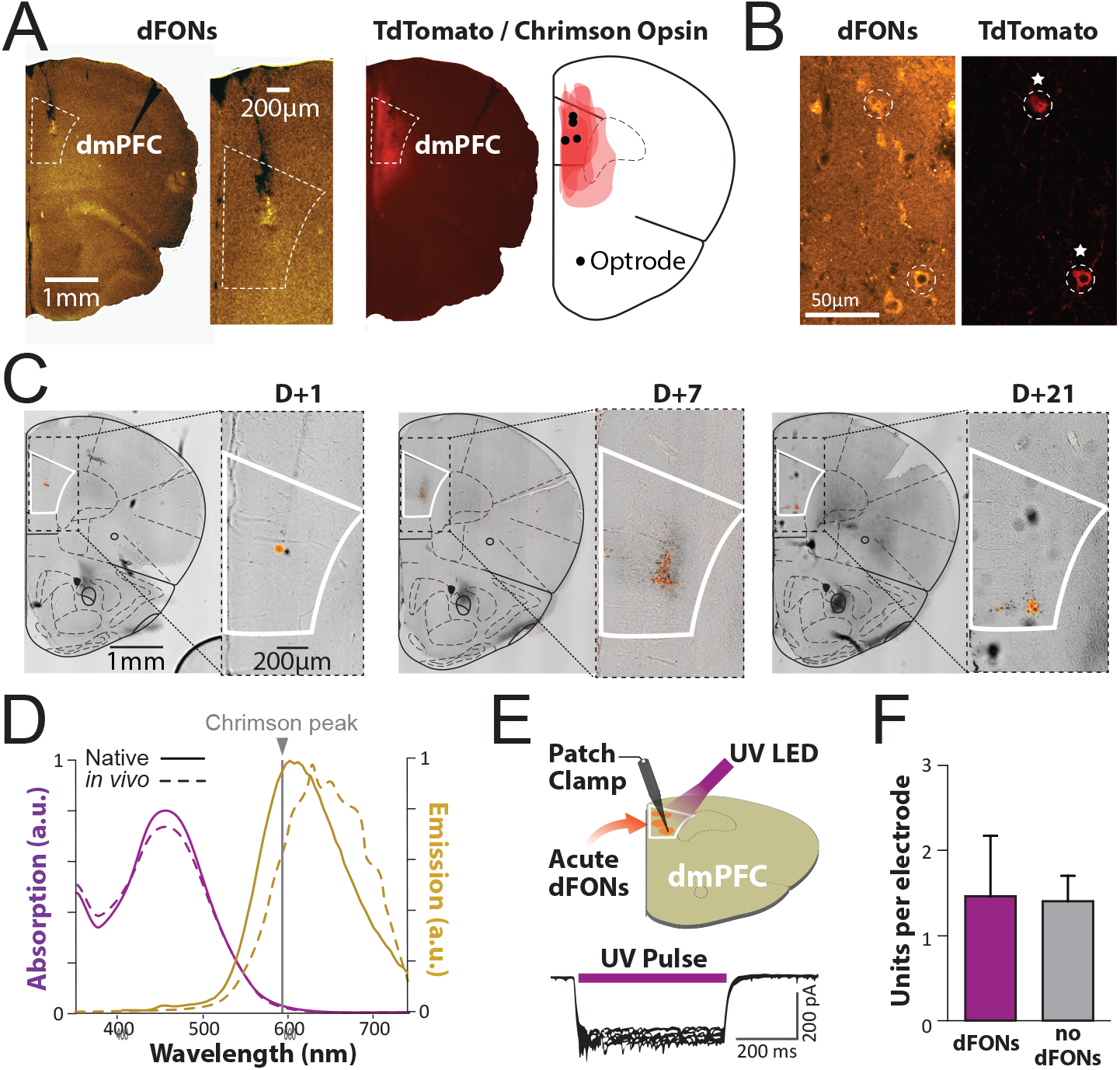
Stability and safety of dFONs **A**@**D** in vivo. A. Brain slices encompassing dorsomedial prefrontal cortex (dmPFC). Left. Example epifluorescence microscopy picture for dFONs **A**@**D** localization in vivo at different magnifications. dFONs deposit is visible at the tip of the optrode track. Right. Corresponding example photograph for viral infection extent and drawing showing infection extent and optrode implantation sites in 4 animals, as well as high magnification parvalvumin interneurons fluorescence. B. Example fluorescence confocal microscopy picture for dFONs **A**@**D** (excitation 405 nm, single plane acquisition 550-650 nm) and tdTomato/Chrimson (excitation 552 nm, single plane acquisition 570-650 nm). Note two transduced PVIN showing bright red fluorescence (dotted circles and star, confirming efficient viral transfection). The colocalisation of dFONs **A**@**D** and tdTomato suggests intracellular cytoplasmic concentration of the nanoparticles. C. dFONs **A**@**D** diffusion remains limited to injection site after their injection. Bright field microscopy pictures for dFONs **A**@**D** diffusion at 1, 7 and 21 days post injection in dmPFC at whole hemisphere and dmPFC magnifcation levels, with fluorescence thresholding in red for dFONs emission. D. Expected photophysical properties of dFONs **A**@**D** are preserved in vivo. Absorption and emission spectra of dFONs **A**@**D** before (plain line) and after injection in dmPFC (dotted line). E. Patch clamp recodring in vitro upon UV light with dFONs **A**@**D** injection. top. Cartoon representing the setup for acute brain slice patch clamp recording neuronal with acute dFONs **A**@**D** application and UV illumination of neurons infected with Chrimson opsin (visualized by orange dots). Bottom. Example of an inward current (depolarizing) during illumination showing that in presence of dFONs **A**@**D**, light pulse elicits a normal response in patched cells expressing the Chrimson opsin. F. Stable neuronal recordings following dFONs. Neuronal activity is maintained after dFONs **A**@**D** injection in vivo. Single unit recordings yield the same amount of neuron with and without dFONs **A**@**D** in dmPFC (t test, t(6)= 0.133, p= 0.899).

*In order to evaluate dFONs* ***A@D*** *efficacy, stability and biocompatibility, the initial phase of this proof of concept aimed at demonstrating: i) the presence of dFONs in close proximity to Chrimson-expressing cells in cerebral tissue over several weeks following injection, ii) the stability of optical properties of dFONs once integrated into brain tissue, and iii) the preservation of physiological neuronal activity in the presence of dFONs*.

#### 2.3.2 In vivo stability of dFONS

The dFONs **A**@**D** were injected in the dorsomedial prefrontal cortex (dmPFC) of Parvalbumin-cre mice previously infected with the virus leading to the selective expression of Chrimson and its fluorescent reporter (tdTomato) in Parvalbumin - expressing interneurons (PVIN, Figure 4A). During the same surgery, we also implanted optrodes - optic fibers bearing recording electrodes-designed to deliver light and record from neuronal activity, respectively. Confocal microscopy imaging of the dmPFC confirmed the presence of FONs up to three weeks post injection, notably at the cellular level, as confirmed by fluorescence in the emission band of dFONs, which could be separated from the emission of the tdTomato, inherent to the expression of Chrimson in PVIN (Figure 4A). To assess the temporal dimension of dFONs diffusion, we harvested brains from animals injected with dFONs at different time points following surgery. Overlay of fluorescence images in the dFONs emission wavelength with transmitted light microscopy showed that they remain concentrated within the limits of dmPFC and close to the injection site at least until 21 days after the surgery (Figure 4B).

To assess the stability of dFONs **A@D** photophysical properties, we measured their absorption and emission spectra on brain slices - around the soma of a neuron- by spectral imaging and compared them to prior-injection, native spectra displayed in Figure 2. In vivo spectra were found to be similar to absorption and emission spectroscopy measures (Figure 2), and the overlap of their fluorescence to the Chrimson absorption peak was adequate for further opsin stimulation (Figure 4C).

We then assessed the impact of dFONs on cell viability. To check that the presence of dFONs did not grossly impair neuronal viability brain slices with dmPFC PV IN expressing Chrimson were harvested for ex vivo patch clamp experiments. The activity of single neurons was recorded while dFONs were applied, and a consistent depolarisation could be observed upon UV light illumination (Figure 4D). Moreover in in vivo electrophysiology experiments with implanted optrodes, the amount of neurons recorded in animals with and without dFONs were similar even after 21 days post injection (Figure 4E). In addition, the fine electrophysiological characteristics of neuronal spiking were found unchanged in all groups of animals (see supporting information, Figure 2). ***All together, these results indicate that dFONs do not critically alter the density or activity of neurons and remain located in close range of the initial site of injection***.

#### 2.3.3 In vivo stimulation of neurons

In order to assess the feasibility of dFONs-mediated optogenetics in vivo, dFONs **A**@**D** were injected into the dmPFC of adult PV-cre mice, resulting in PVIN expressing Chrimson, and also implanted an optrode at the same location (Figure 5A). As illustrated in Figure 5B, PVIN have the property to inhibit neighboring dmPFC principal neurons (PN). Therefore we expect an overall local reduction in the number of PN action potentials upon PVIN optogenetic stimulation. The activity of neurons was recorded under 585 nm or 405 nm illumination - corresponding to the selective excitation of Chrimson and dFONs **A@D** respectively-at various powers (Figure 5). The expected effect of direct excitation (i.e. 585 nm) is an instantaneous activation of PVIN, and a concurrent inhibition of PN, proportional to the intensity of the light. On the other hand, UV illumination is not expected to produce any effect on neurons, unless it is in the presence of dFONs (emitting yellow light) as exemplified in Figure 5C.

**Figure 5:**
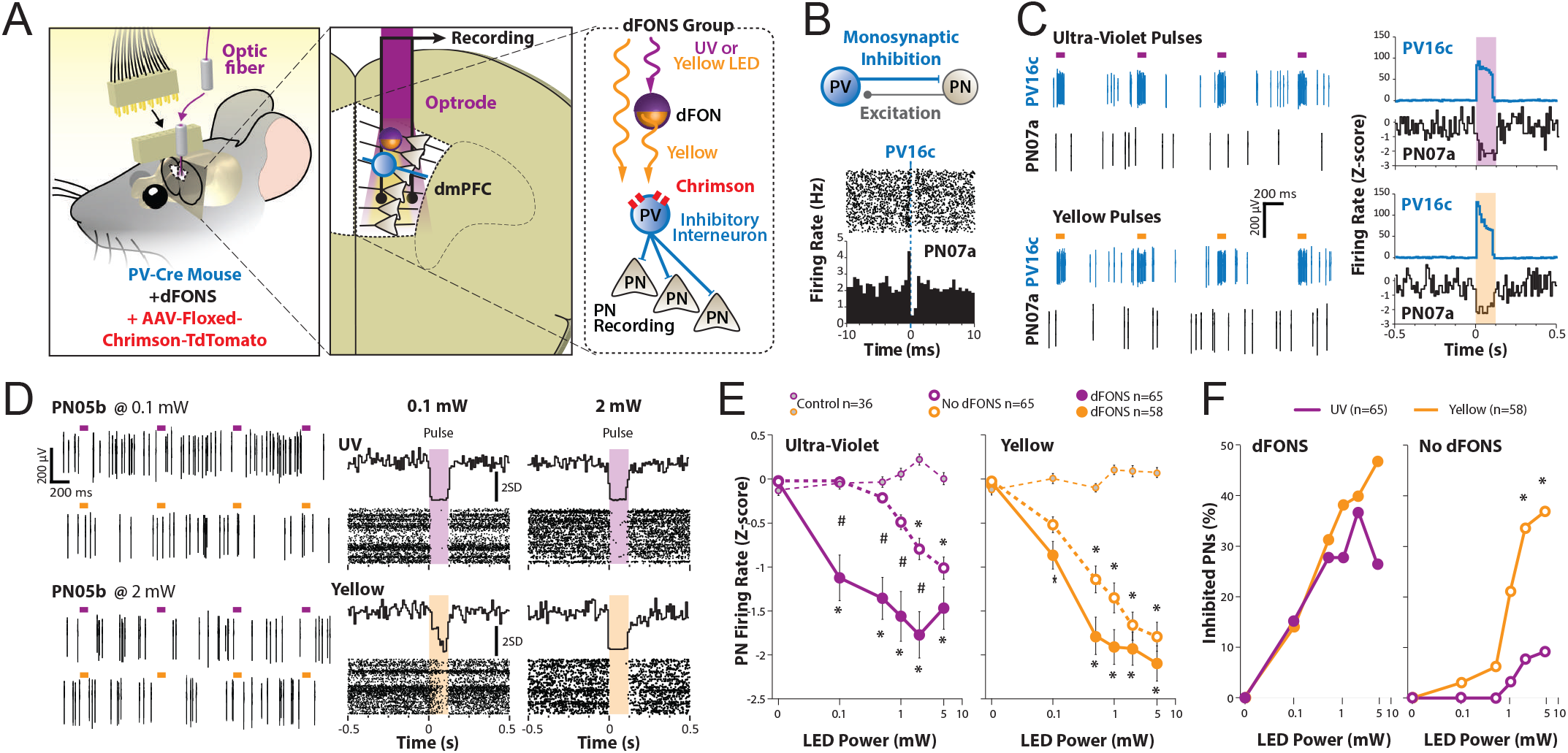
dFONS **A**@**D** efficacy in vivo. A. Cartoon of the experimental setup at different magnification. B. Example of a monosynaptic inhibitory connection between a putative PVIN (PV16c) and a putative principal neuron (PN07a). Note that the presence of a peak before PV16c firing at t=0 indicates that PN07a also excites PV16c reciprocally. C. Left. Raster plots for PV16c and PN07a during UV (top) and yellow light stimulation (bottom). Right. Peristimulus time histogram for PV16c and PN07a centered on light pulse onset for UV (top) and yellow illumination (bottom). D. Left. Example raster plots for a principal neuron (PN05b) under different light color and intensity (top: 0.1 mW; bottom: 2 mW). Right. corresponding peristimulus time histogram for PN05b centered on light pulse onset for UV (top) and yellow illumination (bottom). E. Normalized average firing rate during light pulses of increasing intensity for UV (left) and yellow illumination (right). (2way ANOVA, UV, F1: treatment, F1(2)=120.9; F2: Power, F2(5)=8.5; F1-2(10) = 5.4, p *<* 0.001; Yellow, F1: treatment, F1(2) = 79.4; F2: Power, F2(5) = 27.9; F1-2(10) = 85.4, p *<* 0.001; Bonferroni *post hoc* test. difference with control, p *<* 0.05. # difference with No dFONs, p *<* 0.05). F. Proportion of inhibited PNs as a function of light intensity in the presence or absence of dFONs **A**@**D** (left and right respectively, Chi-Square and Bonferroni multiple comparison with adjusted alpha=0.001, difference between yellow and UV).

In the dFONs group, we observed a significant inhibition of PNs during both direct excitation at 585 nm and indirect excitation at 405 nm (Figure 5D). Analyzing increasing light intensities, effect at 405 nm was observed with power as low as 100 µW while it required 50 to 100 times more power to activate Chrimson without dFONS. A double control (no expression of Chrimson and absence of dFONs) showed unchanged neuronal activity at any light intensity and for both colors. In addition, dFONs presence also lowered the threshold for direct activation at 585 nm as PN firing was significantly reduced at 0.1 mW (0.5 mW without FONs, Figure 5E) and more neurons were impacted at 0.1 and 0.5 mW compared to the condition without dFONs (Figure 5E-F).

Inhibition of PNs upon UV light suggests that i) dFONs-mediated optogenetics allow for a controlled activation of PVIN from UV light, and that ii) dFONs potentiate optogenetic activation of PVIN from yellow light (i.e. 585 nm). The latter observation is attributed to the excitation of the dFONs’ core (via direct absorption of **A**), which is sufficiently bright to activate close-by opsins. This result underscores the potency of dFONs as tools to enhance the efficiency of opsin activation, providing access to lower irradiation thresholds for neuronal manipulations.

#### 2.3.4 In vivo close-loop stimulation

After showing the efficacy of dFONs **A@D** mediated optogenetics on single neurons, we have tested its potency at the functional level by attempting to manipulate animals behaviour, more precisely that of fear expression. We have previously demonstrated that during fear expression, dmPFC activity oscillate at a rhythm of 4 Hz[30] and that PN activity is locked onto the ascending phase of this rhythm [5]. In this study also, we were able to counteract fear by applying a close loop inhibition of PNs in the ascending phase of 4 Hz, by means of a optogenetic activation PVIN, such as the one described in the previous section. The goal here was to reproduce this optogenetic experiment but this time with the mediation of dFONs **A@D**. To this end, we submitted mice to a classical model of Pavlovian fear conditioning (Figure 6A). First, PV-cre mice were injected in dmPFC with dFONs (or saline, no dFONs control group), a virus carrying Chrimson and tdTomato and were bilaterally implanted with optrodes (Figure 6B). For the fear conditioning paradigm, these animals learned to associate a previously neutral auditory stimulus (positive conditioned stimulus, CS+) with a mild electric foot shock (aversive unconditioned stimulus, US). A second neutral stimulus is presented but not associated with any outcome and serves as an internal control (negative conditioned stimulus, CS-). On test day, when exposed to the CS+ alone, animals display a fear response in the form of freezing, which is a temporary cessation of all motor activity but respiration. In animals that properly learned the CS+-US association, freezing is found to be typically increased for the exposition to the CS+, compared to CS-. In this context, the close-loop stimulation consists in the online monitoring of both dmPFC brain waves and animal behaviour and as soon as we detect freezing and 4 Hz ascending phase, we apply UV light (405 nm) for the optogenetic activation of PVIN (Figure 6D), leading in turn to the global inhibition of PNs as shown in Figure 5).

**Figure 6:**
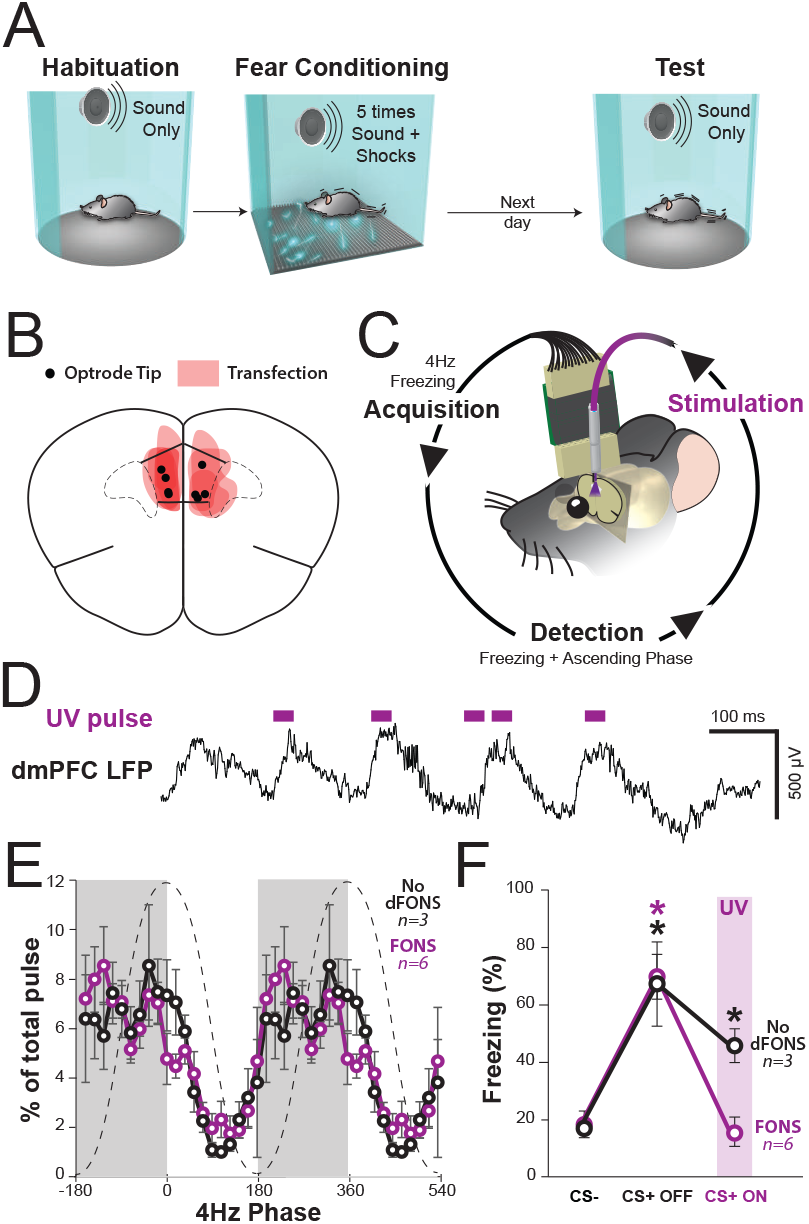
Optogenetic control of fear behavior through excitation of Chrimson opsin from dFONs **A**@**D**. A. Cartoon depicting the protocol of pavlovian fear conditiong and test. B. Drawing of tdTomato-chrimson expression extent and optrode tip locations in all animals in the test group. C. Cartoon representing the principle of close-loop stimulation for realtime inhibition of dmPFC, targeting fear biomarkers: the ascending phase of 4 Hz oscillations in dmPFC and freezing. D. Real-time phase specific stimulation. Example trace showing dmPFC LFP and concurrent UV pulses delivered in the ascending phase of the ongoing 4 Hz oscillation. E. Phase histogram showing light pulse distribution along 4Hz phase for the dFONs **A**@**D** (purple line) and control group (black line, 2way ANOVA, F1: phase, F1(19)=9.6, p *<* 0.001.; F2: treatment, F2(1)=0.0, p = 1; F1x2(19) = 0.7, p = 0.8). E. Effect of ascending phase stimulation on fear behaviour. Histogram of freezing level during conditioned stimuli (CS) presentations. dFONs and control group show a significant increase in freezing level between CS-(neutral) and CS+ (fear conditioned), indicating successful learning. Phase specific stimulation during CS+ (CS+ ON) brings freezing level back to neutral level in the FONs group while no effect was observed in the no dFONs control group which remains elevated (2way ANOVA; F1: treatment, F1(1)=1.1, p = 0.3.; F2: epoch, F2(2)=15.0, p *<* 0.001; F1x2(2) = 3.8, p *<* 0.15; Bonferroni *post hoc* test. difference from CS-, p *<* 0.05).

Behavioural results show that the percentage of time spent freezing was significantly higher for the first block of CS+ than for the block of CS-(CS+ OFF, Figure 6F) indicating that both test (dFONs) and control animals (no dFONs) were effectively conditioned. A second series of CS+ was subsenquently played, but this time they were accompanied by close-loop stimulation with UV light (CS+ ON). Pulse/phase analysis shows that light pulses successfully targeted the ascending phase of dmPFC 4 Hz brain waves in both FONs and no FONs groups (Figure 6D). In the dFONs group this phase specfifc inhibition of dmPFC resulted in a reduction of freezing to a level comparable to that of CS-, while it remained elevated in the control group (Figure 6E). ***Beyond standard optogenetics, this experiment demonstrate the ability of dFONs A@D to mediate behavioural manipulation and therefore be part of a comprehensive, functional neurostimulation toolbox***.

## 3 Conclusion

We successfully prepared core-shell dFONs **A**@**D** by incorporating dye **A** in the core and dye **D** in the shell. Our study establishes that these molecular-based fluorescent nanoparticles exhibit efficient energy transfer from **D** to **A**. Plus, thanks to the NIEE effect, dFONs **A**@**D** have enhanced brightness compared to pure core dFONs **A** under both visible and near-infrared excitations. Remarkably, these nanoparticles exhibit inherent stability in water and in vivo, without requiring any adjunction of polymers nor exogenic surfactants.

We think that the long term stability of dFONs **A@D** within the brain reflects the intrinsic nature of nanoparticles surface: we previously reported that dFONs made of **D** show *stealth* behaviour (i.e. do not have unspecific interactions with cell membranes) when incubated with Hela cancer cells[31]. Such ability of **D** to escape from unspecific interactions was attributed to the electrostatic interactions from polarizable dyes on surface of dFONs with serum proteins, thus forming a stable corona surrounding the nanoparticles. Ritz & al.[32] have studied the effects of protein corona on the uptake of nanoparticles and showed that the nature and the charge of surfactants used for polystyrene nanoparticles strongly influence the type of proteins composing the corona. Other research groups also addressed the influence of the surface chemistry and size of nanoparticles (organic, inorganic and metal-based) on the composition of proteins corona[33, 34]. We hypothesize that the presence of **D** dyes on the outer layer of dFONs **A**@**D** may contribute to the stability of these core-shell dFONs in situ thanks to the presence of a protein corona, limiting their uptake and degradation by cells.

The strong brightness of these dFONs can be exploited for in vivo optogenetics experiments, enabling the activation of opsins with lower irradiation power thresholds compared to the direct excitation of opsins. This demonstrates that dFONs are bright enough to allow substantial activation of opsins despite using wavelengths strongly attenuated in the brain and injecting extremely diluted suspensions (concentration of injected dFONs ~ nM range). ***This paves the route to a better and more efficient activation of Chrimson exploiting the ability of this dFONs to absorbs pulsed NIR light: the large two-photon brightness (****σ*_2_Φ *>* ***2×10***^5^ ***GM) of these nanoparticles in the NIR-I range (from 0*.*70 µm to*** *>* ***1*.*00 µm) promises deeper excitation with potential transcranial neuron stimulation*** [35].

Finally our findings demonstrate the feasibility of using dFONs in vivo for close-loop stimulation, with evidence of efficacy at the single cell (patch clamp), the network (in vivo electrophysiology) and the whole organism (behaviour) levels. Yet, from a clinical standpoint, the use of UV light for 1P stimulation is far from optimal since it requires the implantation of intracranial optic fibers. While UV and visible wavelengths are blocked or diffracted by biological tissues (skin, bone, white and gray matter) NIR wavelength present a much more favorable penetration profile. They could be used for extracranial, non invasive stimulation in combination with NIR to visible photoconverting nanoparticle. Recent studies have successfully achieved this in rodents [9, 10] but with inorganic compounds that are limited in colloidal stability, brightness and toxicity. All of the former are pre-requisites properties for clinical application consideration, and the properties that we managed to confer to **A**@**D** dFONs. An ideal system would combine the safety and stability of the present organic approach with NIR to visible photoconversion properties.

***To the best of our knowledge, it’s the first time that surfactant-, metal- and polymer-free fluorescent organic nanoparticles are used for optogenetic manipulation of neurons and behaviour***. Our results underline that pure dye nanoparticles are promising nano-tools for the enhancement and advancement of optogenetic applications.

## 4 Experimental Section

### General informations

Commercially available compounds (purchased from Aldrich, TCI, Fluorochem or Alfa Aesar) were used without further purifications. DMF used for the Vilsmeier-Haack reaction was distilled and stored with activated molecular sieves (3°A) before use. Spectroscopic grade tetrahydrofuran (THF) was used for the stocks solutions for dFONs preparation. Sonicator probe used for the preparation of dFONs was an ultrasonic processor (model GE 130), and the microsyringes used for the nanoprecipitation were 250 µL and 150 µL Hamilton syringes.

### Synthesis of dyes A and D

Dyes **D** and **A** were synthesized according to the synthesis described in our previous work[23].

### Preparation of dye based fluorescent organic nanoparticles (dFONs)

dFONs **A** or **D** are prepared using the so-called nanoprecipitation method[29]. In a 30 mL vial, 200 µL of a Stock solution of dye **A** (1 mm) or **D** (0.5 mm) in THF was quickly added in 20 mL of distilled water under sonication (10 W output). Instantaneously, the solution turns coloured and the sonication was left for 3 min. The clear and coloured solution was then allowed to cool down to room temperature, then used without further purifications.

### Preparation of core-shell dye based fluorescent organic nanoparticles (dFONs A@D)

dFONs **A**@**D** are prepared by a two-step procedure: In a 30 mL vial, 200 µL of stock solution of dye **A** (1 mm in THF) was quickly added in 20 mL of distilled water under sonication (10 W output) for 3 min. The clear and reddish solution was left to cool down at room temperature. 10 mL of the dFONs **A** solution were collected and 100 µL of a stock solution of dye **D** (0.5 mm in THF) were added dropwise (1 drop / 5 sec) with strong magnetic stirring (600-800 rpm) for 10 min.

### Electronic microscopy

Size of the dFONs **A** and dFONs **A**@**D** were measured using transmission electronic microscopy (TEM) at Bordeaux Imaging Center (BIC) facility. TEM images were acquired on a Hitachi H7650 at 80 kV. dFONs were deposited on copper grid previously charged (+) by Glow discharge technique. A drop of dFONs was left on the charged grid for 1 min., and the excess of liquid was gently removed with a paper by capillarity. Then, a drop of aqueous uranyl acetate was added on the dFONs stained grid for 3 min. Afterward, the excess of liquid was gently removed with a paper by capillarity. The size distribution was determined by manual counting of dFONs (N= 517 for dFONs **A** and N= 434 for dFONs **A**@**D**) using ImageJ software. A binning of 5 was used for the generation of histograms.

### Photophysical properties of dFONs

All linear and non-linear photophysical properties were measured at NanoMultiPhot platform (ISM, University of Bordeaux). Absorption measurements were performed on V670 UV-Vis spectrophotometer (JASCO) at room temperature with 1 cm quartz cuvettes. Fluorescence measurement were conducted on diluted solution (as such the maximum of absorbance was in the range 0.05-0.10) using a FluoroLog3 (Horiba) and 1 cm (four faces) quartz cuvettes. Fluorescence quantum yields (Φ_*f*_) were calculated using equation 1.

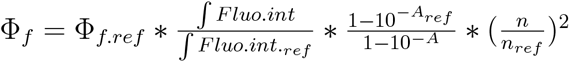

Equation 1. Calculation of fluorescence quantum yield Φ_*f*_

Where, ∫*Fluo*.*int*. is the integral of the fluorescence intensity, A is the absorbance at excitation wavelength, n is the refractive index of the solvent and ref the standard. Fluorescence lifetimes measurements were done on a FLS1000 (Edinburgh Instruments) using a NanoLed-370 with Controler NL-C2 (both Horiba) and an EPL-510 (Edinburgh Instruments) for excitation at 370 nm and 504 nm respectively. Aqueous solution of Ludox (30% in water, Sigma-Aldrich) was used for instrument correction and deconvolution.

Two-photon absorption measurements were performed on aqueous dFONs suspensions (C = 10 µm for dFONs **A, A**@**D** and 5 µm for dFONs **D**) using fluorescein in aqueous NaOH (C = 10 µM, pH 11.0)[36, 37] as two-photon reference. The two-photon absorption was measured using the well-established method described by Xu and Webb[37]; briefly two-photon absorptions were measured using the Two-Photon Excited Fluorescence technic (i.e. TPEF). To do so, an Ultra II (Coherent) femtoseconds pulsed laser (repetition rate: 80 MHz, pulsed width: 150 fs) was focalised on dFONs or fluorescein solutions using an air Olympus objective (10x, 0.25 NA) under magnetic stirring. Excitation wavelength was tuned in the range 700-1000 nm. The 1 cm quartz cuvette was set in order to have the fluorescence generated on the close vicinity of the wall of the cuvette to limits the reabsorption of fluorescence. The fluorescence is then collected in epifluorescence and filtered from back-scattered laser using a dichroic mirror (675dcxru, Chroma) and finally send to a fast spectrophotometer (MayaPro, OceanInsight) by the mean of an optical fiber. The power of the laser is tuned using a polarizer at Brewster’s angle (142013, Layertec) in tandem with a rotating halfwave plate. The power is measured in real time by sending a part of the laser beam on a silicon photodiode (S142CL with power meter interface PM101U, both from Thorlabs) by the means of a beam splitter (UFBS5050, Thorlabs). Quadraticity of the TPEF signals is measured for each excitation wavelength checking the quadratic dependence of fluorescence to the laser power (data available in the supporting information, Figure 1). The full programs used for TPEF measurements and analysis are available on github[38]. The two-photon brightness (*σ*_2_Φ) of the dFONs was calculated using equation 2.

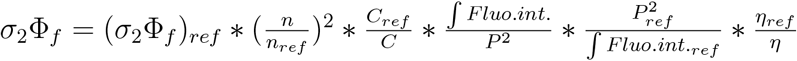

Equation 2. Calculation of two-photon brightness, σ_2_Φ_*f*_.

Where n is the refractive index of the solvent, C the concentration, P the laser power, J *Fluo*.*int*. the fluorescence intensity and *η* the spectral correction of the dFONs and reference (i.e. ref). For dFONs **A@D** the concentration used for the calculation of *σ*_2_Φ_*f*_ was set to the concentration of the emitter (i.e. dye A).

### Origin, housing and ethical approbation for animal experimentation

Wild type male and female C57BL6/J (11-18 weeks old, Janvier), and Pva-cre (11-18 weeks old, Janvier) were housed in rooms of controlled temperature and humidity with a 12 h light-dark cycle and provided with food and water ad libitum. Mice were housed individually starting from the day of the first surgery. All procedures were performed per standard ethical guidelines (European Communities Directive 86/60-EEC) and were approved by the Animal Health and Care committee of Institut National de la Santé et de la Recherche Médicale and French Ministry of Agriculture and Forestry (agreement A33-063-099).

### Surgery

Pva-cre mice underwent one or two surgeries (1. viral infection; 2. dFONS injection and optrode implantation) while wild type animals (age 8-11 weeks) were only subjected to dFONs injection. For the viral infection surgical procedure, Pva-cre were anesthetized with isoflurane (Vetflurane®, from Virbac, induction 5%, maintenance 1.5%) in air. Body temperature was maintained at 37°C with a temperature controller system (FHC) and eyes were hydrated with an ocular moisturizing gel (Ocry-gel, from TVMlab). Mice were secured in a stereotaxic frame (Kopf Instruments) and had their skin shaved, cleaned and disinfected with povidone-iodine (Betadine). Meloxicam (Metacam® from Boehringer Ingelheim, 4 mg/kg in 0.2 mL) was injected subcutaneously in the back of the mice, and lidocaine (Lidor® from Axience, 40 mg/kg in 0.1 mL). Both were diluted to their target concentration with sterile water (CDM Lavoisier). We injected a cre-dependent adeno associated virus carrying (AAV-flex-Chrimson-tdTomato) loaded in glass pipettes. These pipettes were pulled from glass capillaries with an end diameter of 40-50 µm were filled with the solution to be injected. The solution was delivered using a custom manual air pumping system or a one-axis hydraulic micromanipulator (Narishige, MO-10). During the surgery, pipettes were lowered into the prefrontal cortex according to coordinates relative to the bregma and brain surface (2.0 mm anterior to the bregma, 0.4 mm lateral to midline and 1.2 to 1.4 mm ventral to the cortical surface), revealed by drilling cortical bone. Once the target coordinates were reached, a waiting time of 5 minutes was respected before injecting the solution at a controlled flow under 60 nL/min. Once the injection complete, the pipette was left in place for 5 additional minutes before slow removal. Viruses were all delivered in a 180 nL volume per target brain area. After this surgery, mice were resting until awake in their heated homecage. Pva-cre mice were allowed to recover under close monitoring for 7 days minimum, and were housed without any further intervention for 3 weeks in order to achieve significant Chrimson/tdTomato expression. A subset of animals then entered the ex-vivo patch clamp experiment track (see below) while a second subset entered in the in vivo track and were also submitted to the following procedure.

The injection and/or implantation surgery was carried under the same anaesthesia and pain management methodology as above. PV-cre and wild type animals both received an prefrontal injection of photoactive nanoparticules in a 1000 nL volume in similar pipettes and with similar coordinates as above. Pva-cre animals were also implanted in prefrontal cortex with an optrode (optic fiber + microelectrodes). This implants was secured using Super-Bond cement (Sun Medical) and polyacrylate glue. After this surgery, mice were resting until awake in their heated homecage. Pva-cre mice were allowed to recover under close monitoring for 7 days minimum, before being habituated to gentle handling and enter further in vivo experimental procedures. Wild type mice would be sacrificed at 1, 7 or 21 day following the surgical procedure (see perfusion and histology section below)

### Optrode fabrication

Optrode consisted in microelectrodes affixed to an fiber optic designed for the recording of neurophysiological signals and light delivery, respectively. Fiber optic implants were either built in lab from zirconium ferrules (Prizmatix) and quartz optic fibers (Thorlabs, NA= 0.5 to accommodate UV transmission) solidarized with epoxy glue, cut at the desired length with a diamond cutter and mechanically polished, or purchased (Prizmatix). All fibers had 1.25 mm diameter ferrules with a 200 µm diameter fiber (core and cladding), the latter protruding at a consistent length from the cannula (usually, at 2 mm + depth of the target brain area from brain surface, about 3-4 mm total for prefrontal cortex fibers). All optic fibers were checked for light transmission using an optical power meter (Thorlabs), and fibers with a transmission out of the 65-85% range were rejected.

The recording electrodes were made of 32 individually insulated nichrome wires (13 µm inner diameter) with a matched impedance of 65-75 kΩ for bundle, and 140-150 kΩ for tetrodes (adjusted by electroplating in a solution of gold nanoparticles and carbon nanotubes) configurations. Tetrodes were built using a custom system involving a magnetic stirrer and a heat gun to join the 4 wires forming it through their fused insulation. The ends of the electrodes were attached to the extremity of a 200 µm diameter optic fiber using polyacrylate glue and sucrose water on the portion meant to enter the brain tissue, to reduce chemical foreign body reaction. The electrodes were protruding from the end of the fiber in order to record from neurons encompassed in the cone of light, that is at distances affected by local optogenetic effects in the literature. The wires were attached to two 18-pin connectors (Omnetics) or a single 36-pin connector (Omnetics).

### Ex vivo patch clamp experiments

Wild type mice were used at the age of 4–9 weeks. The extracellular artificial cerebro-spinal fluid (ACSF) solution utilized for slice recordings is composed of: 119 mm NaCl; 2.5 mm KCl; 1 mm MgCl2; 2 mm CaCl2; 10 mm glucose; 1 mm NaH2PO4; 26 mm NaHCO3. The cutting solution is an ice-cold sucrose solution (1–4 °C) composed of 2 mm KCl; 2.6 mm NaHCO3; 1.15 mm NaH2PO4; 10 mm glucose; 120 mm sucrose; 0.2 mm CaCl2 and 6 mm MgCl2. Both solutions are oxygenated with carbogen (95% O2, 5% CO2, pH 7.4 at 37 °C, 290–310 mOsm/L). For brain dissection, mice are anesthetized with 5% isoflurane for 2 min before decapitation. The head is immersed in the iced sucrose solution. The removed brain is immersed for 4 min in iced oxygenated sucrose solution and then placed on a cellulose nitrate membrane to separate and position the hemispheres in the vibratome (Leica VT1200s) to obtain 320 µm coronal slices (cutting speed of 0.12 mm/s). Once produced, slices are semi-immersed in a dedicated incubation chamber, oxygenated and maintained at 35 °C for at least 2 h before starting the recordings. Recordings are made in an S-shaped recording chamber, maximizing oxygenation while preventing slice movement caused by the 2 mL/min perfusion flow. Whole cell patch clamp recordings are obtained using borosilicate glass micropipettes (4 to 6 megaohms) stretched with a PC-10 (Narishige, Japan). The pipette is filled with an intracellular solution containing (in mm): 125 CsMeSO3, 2 MgCl2, 1 CaCl2, 10 EGTA, 10 Hepes, 4 Na2-ATP, 0,4 Na-GTP. Electrophysiological recordings are obtained using a MultiClamp 700B (Molecular Devices, Foster City, CA) with Clampex 10.7 software (Molecular Devices, Foster City, CA).

A second pipette was loaded with dFONs solution in order to add the nanoparticles to the slice and test their influence on UV light stimulation of Chrimson.

### Light sources and delivery

Implanted cannulae were adapted to a simple or two ends optic fiber for single hemisphere or dual hemisphere light delivery (Doric/Thorlabs, or Prizmatix) through a zirconium mating sleeve (Prizmatix). This fiber would freely hang above the animal during all of the experiment, and was connected to a patch cord fiber (Thorlabs or Prizmatix) through a rotary joint (Thorlabs or Prizmatix). This patch cord fiber was finally linked to the light source, which was either a LASER (473 nm [blue], 532nm [green]) or an LED (Dual-FC-LED 405 nm max, [UV], 460 nm max, [blue] 545 nm max; FC-LED-green 585 nm max; FC-LED-yellow 655 nm max; FC-LED-deep red, Prizmatix). A pulse generator (Master 9, AMPI) was parametered to generate direct current to control the activation of the LASER or diode light source through a BNC coaxial cable. Generator pulse currents were collected on the recording setup to keep track of the beginning and end of stimulation timestamps. For PSTH activity modulation depending on injected power and color, the stimulation was set to 100 ms duration at 1Hz.

### Fear conditioning paradigm and behavioural analysis

Auditory fear conditioning and testing took place in two different contexts (context A and B). The conditioning and testing boxes (IMETRONIC) were cleaned with 70% ethanol and 1% acetic acid before and after each session, respectively. On day 1, mice were submitted to a habituation session in context A, in which they received four presentations of the CS+ and of the CS-(total CS duration, 30 s; consisting of 50-ms pips at 0.9 Hz repeated 27 times, 2 ms rise and fall, pip frequency, 7.5 kHz or white-noise, 80 dB sound pressure level). Discriminative fear conditioning was performed on the same day by pairing the CS+ with a US (1 s foot-shock, 0.6 mA, 5 CS+-US pairings, inter-trial intervals, 20-180 s). The onset of the US coincided with the offset of the CS+. The CS-was presented after each CS+–US association but was never reinforced (five CS-presentations; inter-trial intervals, 20-180 s). On day 2, conditioned mice were submitted to a testing session (Retrieval session) in context B during which they received 4 and 12 presentations of the CS- and CS+, respectively, while being filmed and having their dmPFC electrophysiological activity recorded (Neuronal System Processor, Blackrock Microsystem). For optogenetic experiments, light stimulation would be activated by the close-loop system in the second 4-CS+ block (From the 5th to the 8th presentation of the CS+ during retrieval), in order to compare the behavior to CS+ presented without light (CS+OFF, 1st to 4th CS+) to those with light (CS+ON, 5th to 8th CS+). All sessions were systematically recorded by an infrared camera (30Hz sampling rate) coupled to a video tracking system returning the instantaneous position of the animal (NeuroMotive, Blackrock Neurotech). From this, we derived the instantaneous speed of the animal. We score freezing behavior with a custom Matlab script designed to quantify the moment to moment amount of locomotion of the animals. Mice were considered to be freezing if no movement was detected for 2 s.

### Electrophysiological recordings

Electrodes connectors were plugged in to two 16-channels or a single 32-channels preamplifiers and digitized at 30 kHz (Blackrock Microsystems). For single neuron spike sorting, signal was bandpass filtered from 250 Hz to 5 kHz, and spikes were detected by time-amplitude window discrimination using the Neuronal Signal Processor (Blackrock Microsystems). For local field potential analysis, signal was bandpass filtered from 0.5 Hz to 250 Hz and downsampled to 1 kHz using the Neuronal Signal Processor (Blackrock Microsystems).

### Single-unit analyses

Single-unit spike sorting was performed using Off-Line Spike Sorter (Plexon) for all behavioral sessions. Principal-component scores were calculated for unsorted waveforms and plotted in a three-dimensional principal-component space. Clusters containing similar valid waveforms were manually defined. A group of waveforms were considered to be generated from a single neuron if the waveforms formed a discrete, isolated cluster in the principal-component space and did not contain a refractory period less than 2 ms, as assessed using auto-correlogram analysis. To avoid analysis of the same neuron recorded on different channels, we computed cross-correlation histograms. If a target neuron presented a peak of activity at a time that the reference neuron fired, only one of the two neurons was considered for further analysis. To separate putative inhibitory interneurons (INs) from putative excitatory principal neurons (PNs) we considered three dimensions: spike half-width, spike area under the curve and firing rate. This three dimensional space was then clustered with a k-means method. Neurons with a short wave form and a high firing rate were classified as putative INs while neurons with a longer wave form and a lower firing rate were classified as putative PNs (see supporting information, Figure 2B). This approach was complemented by optogenetic phototagging with PNs having to present an absence of excitation or an inhibition at the time of PVINs light driven excitation.

### LFP 4Hz phase analysis

Local field potentials were analyzed using custom-written Matlab programs. For phase analysis, the signal was filtered in the 2-6 Hz frequency band (4 Hz brain waves) and the complex-valued analytic signal was calculated using the Hilbert transform.

The vector length and the arctangent of the vector angle provide the estimation of the instantaneous phase of the signal, respectively, at every time point. A phase of 0° corresponds to the peak of prefrontal waves.

### Close-loop stimulation

To apply our stimulation in a functionally specific manner, we performed optogenetic inhibition of PNs as a function of the phase (ascending or descending) of the ongoing 4 Hz oscillation in dmPFC LTP. To that end we designed a close-loop stimulation protocol where animal behavior (freezing) and dmPFC LFP (4 Hz power and phase) were monitored online and simultaneously analyzed to drive the stimulation. Animal position was sampled at 30 Hz and LFP at 1 KHz, then uploaded every 30 ms in Matlab for online analysis. Thus analysis consisted in 1/ quantifying animal speed in the last 2000 ms and processing the LFP signal to retrieve instantaneous 4 Hz power and phase. Animal speed was calculated as the average distance travelled over the last 2000 ms time window and compared with the freezing threshold. Bandpass filtering was applied using a second-order Butterworth filter and Hilbert transform was used to estimate both 4 Hz power in the last 500 ms and LFP instantaneous phase for the last retrieved data point. From there, three criterias were defined for the most recent 30 ms upload in order to trigger the stimulation the stimulation. 1/ animal speed was below freezing threshold for at least 2000 ms, 2/ 4 Hz power was above that of baseline level and 3/ 4 Hz phase was encompassed within the range of choice (0-180°). Shall the three criteria be met, then stimulation was triggered. The stimulation consisted of a 1ms pulse sent from Matlab to a pulse generator (Master 9, AMPI) that in turn sent a 30 ms pulse to the LASER or diode generating the light for optical stimulation to the dmPFC. The whole computation from neuronal data read to LASER onset was achieved in 30 ms maximum (data retrieval 10 ms, computation 5-20 ms). Hence, at the end of the 30 ms light pulse, a new analysis loop was completed on the most recent 30 ms upload and shall the three criteria be met then a new stimulation pulse was triggered.

### Perfusion and histological sample preparation

All Mice were administered a lethal dose of pentobarbital with lidocaine (Exagon® from Richter Pharma, 30 mg/kg, and Lidor® from Axience, 30 mg/kg, in 0.3 mL) via intraperitoneal injection and underwent transcardiac perfusion in the left ventricle of 4% paraformaldehyde (PFA, 4% mass/volume) in phosphate buffer saline (PBS, 0.1 m). In electrode-implanted Pva-cre animals, electrolytic lesions of 10 µA were applied for 5 s using a high current stimulus isolator (World Precision Instruments) to identify electrode bundle tip location prior to PFA perfusion. Following dissection, brains were post-fixed for 24 h at 4 °C in 4% PFA before being transferred to PBS with sodium azide buffer (0.02% mass per volume), and kept at 4 °C until histological processing. Brain sections of 50 or 70 µm anteroposterior thickness (for immunohistochemistry or implantation and viral infection verification alone, respectively) were cut on a vibratome, mounted on microscope slides with VectaShield DAPI-loaded mounting medium (Vector Laboratories) and imaged using an epifluorescence system (Leica DM 5000) or a confocal microscope.

### Brain sections imaging

Images were acquired using a Leica SP8 confocal microscope operated with LAS X software, through a 40x NA 1.3 oil objective (Leica HCPL APOCS2). Pinhole was set to around 1 airy and adjusted to preserve a constant optical slice thickness across channels (1.04 µm). FONs were visualized using a 405 nm emitting laser for excitation and a hybrid detector with a 550-650 nm acquisition window. tdTomato (fused with Chrimson) was visualized using a 552 nm emitting laser for excitation and a PMT with a 570-650 nm acquisition window. Cross-talk was prevented by sequential acquisition of the two channels and by using a 405 nm excitation which is in theory out of the tdTomato excitation spectrum. Only linear adjustments were applied to the entire images for the creation of figures.

### Statistical analyses

For each statistical analysis provided in the manuscript, the Kolmogorov-Smirnov normality test was first performed on the data to determine whether parametric or non-parametric tests were required. Two different approaches were used to calculate the sample size. For studies in which we had sufficient information on response variables and expected effect, as PSTH and behavior, power analyses were performed to determine the number of mice needed. For studies in which the effect of the variables could not be pre-specified, as cell counting in histology, we used a sequential stopping rule. In essence this method enables null-hypothesis tests to be used in sequential stages, by analyzing the data at several experimental points using the appropriate tests. Usually the experiment started by testing only a few animals until a mechanistically relevant parameter was significant. This parameter was then pre-defined and incorporated in the null hypothesis for a significant effect, and applied to replicates of the same experiments with appropriate controls. Sample size determination using sequential stopping rule analyses were used for histological counts analyzes in which we could not predict the effect based on previous experiments completed in the team. Using this strategy, we ended up with a value of n comprising between 4 and 5 animals per group. The groups were blinded and shuffled virus-wise (opsin vs reporter alone) and compound-wise (nanoparticles vs PBS alone).

## Supporting information

Supporting informations

## Supporting Information

Supporting Information is available from the Wiley Online Library or from the author.

## Acknowledgements

This work was supported by the French National Research Agency (program NEONS, ANR-22-CE18-0035-02). J.L. and E.K. acknowledge the financial support of University of Bordeaux for their PhD grants. Electronic microscopy was performed at the Bordeaux Imaging Center, a service unit of the CNRS-INSERM and Bordeaux University, member of the national infrastructure France BioImaging supported by the French National Research Agency (ANR-10-INBS-04).

