## Supporting informations for "Dye-based Fluorescent Organic Nanoparticles, New Promising Tools for Optogenetics"

F.L.
Université de Bordeaux, Institut Interdisciplinaire des Neurosciences, CNRS UMR5297, Bordeaux, France

E. K., J.-B. V., M. B.-D., J.D.
Université de Bordeaux, Institut des Sciences Moléculaires, UMR CNRS 5255, Talence, France

### 1 Two-photon absorption

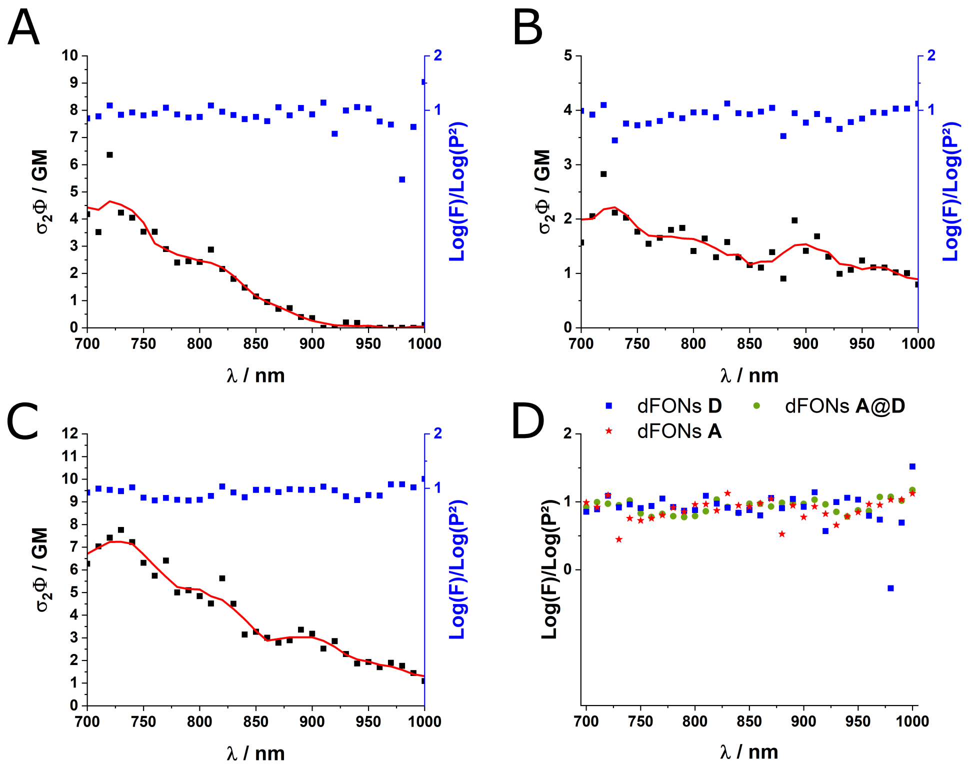


Figure 1: Validation of two-photon absorption processes. Verification of the quadraticity of the fluorescence to the incident power of the laser for dFONs **D** (A), **A** (B) and **A@D** (C) for each excitation wavelengths, overlay with the measured two-photon brightness of the corresponding dye(s) in water. An overlay of the check of the quadraticity of the signal for the three dFONs is shown in D.

### 2 Electrophysiology

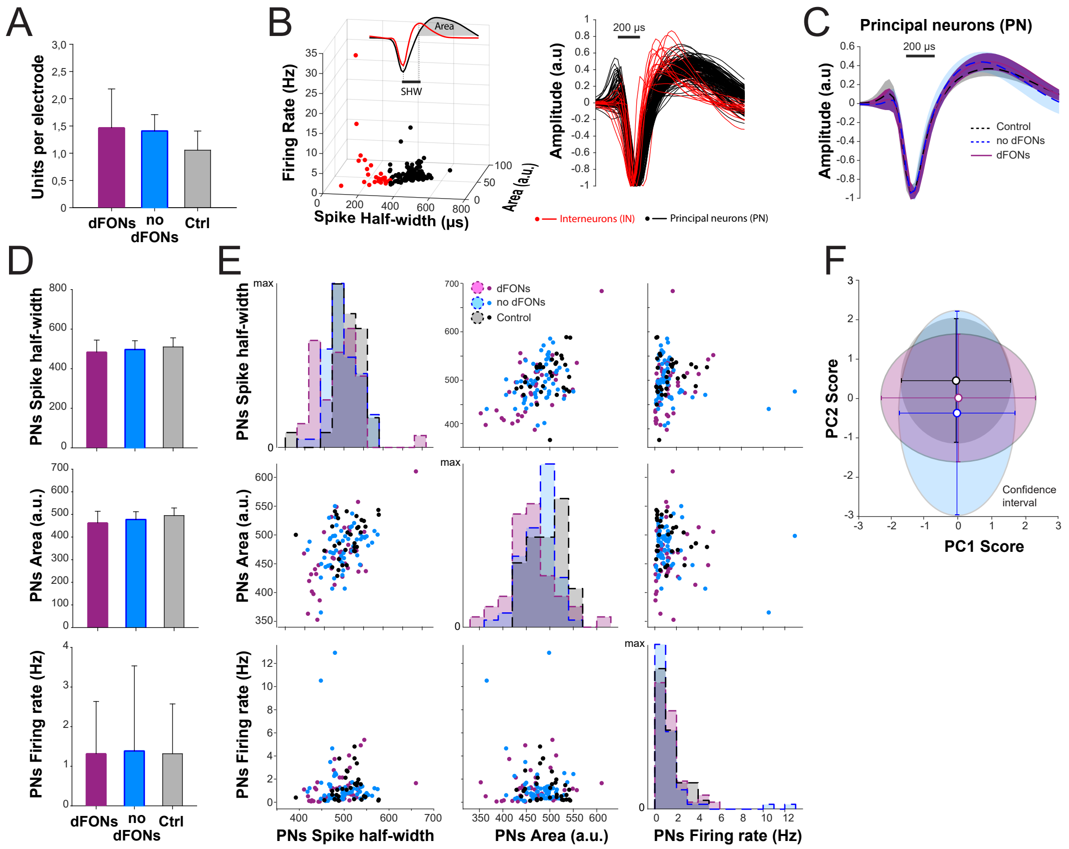


Figure 2: Spike statistics are preserved in the presence of dFONs **A@D** A. The amounts of single neurons discriminated with our without dFONs are similar (t-test, p¡0.05). B. Separation of cell types. For cell type segregation we considered three statistical dimensions of spikes : spike half-width, spike area under the curve and firing rate. K-means clustering returned a group of neurons with a short wave form and a high firing rate (putative inhibitory interneurons: INs, red) and a group with a longer wave form and a lower firing rate (putative excitatory principal neurons: PNs, black) C. Average PN waveforms in dFONs, no dFONs and control groups strongly overlap. D. Average spike half-width (top), spike area under the curve (middle) and firing rates (bottom) for the three experimental groups. E. Diagonal: Superimposed distribution of spike half-width (top), spike area under the curve (middle) and firing rates (bottom) for each group. outside diagonal: two dimensional space representing each dimension against another. One point point is a spike recorded in one of the groups (purple: dFONs, blue: no dFONs, black: control). The 2D distribution of each group strongly overlap. E. Spike shape statistics do not differ from one group to another. In order to formally compare the spike statistics between each group, we applied dimensionality reduction to our data by means of principal component analysis (PCA). PC score distributions did not differ from one group to the next (MANOVA, $\lambda$:0.9129,0.9995, p=0.97).
